# 3D neuroblastoma models expose divergent responses to magnetic hyperthermia and photothermal therapy

**DOI:** 10.64898/2026.07.06.736677

**Authors:** Gema Quiñonero, Andreia P. Magalhães, Lara Diego-González, Juan Gallo, Jaume Mora, Josep Samitier, Aranzazu Villasante

## Abstract

**Purpose:** Hyperthermia is emerging as an adjunct strategy in pediatric oncology, yet its translation is limited by poor understanding of how different modalities impact complex tumor microenvironments. Neuroblastoma (NB), the most common extracranial solid tumor in children, displays profound heterogeneity that hampers therapeutic predictability. Here, we performed the first systematic head-to-head comparison of photothermal therapy (PTT) and magnetic hyperthermia (MH) in tissue-engineered NB (TE-NB) models.

**Methods:** TE-NB scaffolds incorporating five NB cell lines were loaded with magnetic nanoparticles (MNPs) and subjected to PTT (808-nm laser, 130 W/cm^2^, 10 min) or MH (285 kHz, 20 mT, 60 min). Constructs were analyzed at 24 h, 48 h, and 5 d post-treatment for DNA content, cell viability, proliferation (Ki67 immunohistochemistry), and apoptosis (caspase-3/7 staining).

**Results:** MH produced consistent MNP-dependent heating with minimal background, while PTT was dominated by nonspecific medium absorption. Both modalities modulated proliferation within 24 h, but effects varied sharply by cell line and scaffold region, reflecting microenvironmental heterogeneity. By 48 h, PTT often triggered paradoxical increases in proliferation, whereas MH disrupted scaffold integrity, reduced DNA content, and suppressed Ki67 expression. Notably, neither modality induced sustained caspase-3/7 activation, indicating that cytotoxicity proceeds via non-apoptotic pathways.

**Conclusion:** Our findings position MH as a superior modality for uniform heating and proliferation control in 3D NB models, but also highlight that hyperthermia should be considered a context-dependent modulator rather than a binary cytotoxic agent. By integrating patient-specific TE-NB platforms, “precision hyperthermia” could define individualized therapeutic windows, optimize combinations with pro-apoptotic or immunomodulatory agents, and accelerate translation of hyperthermia strategies for children with NB.

## Introduction

Neuroblastoma (NB) is the most common extracranial solid tumor of childhood and a leading cause of pediatric cancer mortality^1^. It typically arises from sympathetic nervous system progenitors (commonly in the adrenal medulla or paraspinal ganglia) and usually presents in infants and toddlers. Indeed, the median age at diagnosis is around 17 months^2,3^. Globally, NB accounts for over 15% of all cancer-related deaths in children. Epidemiological data show that its incidence and mortality have increased over the past few decades, particularly in regions with limited medical resources^1^. Recent Global Burden of Disease analyses predict that without new interventions, the worldwide incidence, mortality, and disability-adjusted life years (DALYs) due to NB will continue to rise through at least 2035^1^. Notably, the etiology of NB remains poorly understood, and no clear modifiable risk factors have been confirmed^4^, making primary prevention difficult and highlighting the need for effective treatments.

NB exhibits marked clinical heterogeneity and is classified into risk groups (low, intermediate, and high-risk) based on features like age, stage, and tumor genetics^5^. Outcomes vary dramatically by risk group: survival exceeds 95% in low-risk NB, whereas long-term survival is only ∼50% in high-risk NB cases^4^. High-risk disease, which represents roughly half of all NB diagnoses, remains therapeutically challenging^5^. Despite intensive multimodal treatment (combining surgery, chemotherapy, stem cell transplant, and immunotherapy), outcomes for high-risk NB have plateaued, with almost half of patients experiencing relapse or progression that proves fatal^5^. This lack of a breakthrough cure for refractory NB highlights an urgent need for innovative therapeutic strategies to improve survival in this aggressive pediatric cancer.

One emerging therapeutic strategy for solid tumors involves hyperthermia-based nanotherapies, notably magnetic hyperthermia (MH) and photothermal therapy (PTT), which eradicate cancer cells through localized heating^6^. In PTT, photothermal agents such as gold nanorods or carbon-based nanostructures absorb near-infrared (NIR) light and convert it into heat via non-radiative relaxation^7^. This provides precise thermal ablation in superficial or accessible tumors. However, light scattering and absorption in deep tissues significantly limit NIR penetration, often necessitating invasive placement of interstitial laser fibers to deliver energy to tumors beyond a few millimeters in depth^8^.

By contrast, MH uses magnetic nanoparticles (MNPs), typically superparamagnetic iron oxide, that dissipate heat under alternating magnetic fields (AMF) through Néel and Brownian relaxation mechanisms^9^. Unlike light, magnetic fields penetrate uniformly into tissues, enabling non-invasive heating even in deep or anatomically complex tumor sites^10^. Importantly, iron oxide nanoparticles are already FDA- and EMA-approved for clinical use in humans, including pediatric patients, as MRI contrast agents and iron supplements, suggesting a favorable safety profile and accelerating translational potential^11,12^ and they have been reported as feasible PTT mediators.

Both PTT and MH are considered minimally invasive and can synergize with chemotherapy, radiotherapy, and immunotherapy by sensitizing tumor cells to treatment, enhancing tumor penetration, and, in some cases, promoting immunogenic cell death^6^. Given their ability to deliver localized cytotoxicity while sparing healthy tissues, and the established safety of certain magnetic nanoparticles in children, these nanotherapy approaches hold particular promise for pediatric cancers, where treatment toxicity and long-term sequelae remain major clinical challenges.

In the context of neuroblastoma, several preclinical studies have demonstrated the feasibility and therapeutic benefit of both PTT and MH. Interstitial PTT using Prussian blue nanoparticles, guided by real-time ultrasound, has been tested in TH-MYCN–driven syngeneic mouse tumors, achieving more uniform intratumoral heating than surface illumination and resulting in greater tumor regression and durable survival^13,14^. Similarly, polysialic acid-imprinted “nanomissiles”^15^ or multifunctional co-delivery platforms combining PTT with differentiation therapy (retinoic acid) and chemotherapy (doxorubicin)^16^ have shown enhanced efficacy in xenograft models by targeting NB heterogeneity and sensitizing tumors to treatment. Magnetic hyperthermia approaches, such as fluorodeoxyglucose-conjugated magnetite nanoparticles, have so far been restricted to in vitro studies with human neuroblastoma cell lines, demonstrating selective uptake and cytotoxic heating under alternating magnetic fields^17^.

Despite these advances, a critical gap remains: no studies have yet evaluated nanoparticle-mediated hyperthermia in advanced three-dimensional (3D) systems of NB. 3D models have become indispensable to better reproduce tumor biology. Conventional 2D monolayer cultures fail to capture NB’s cellular heterogeneity, spatial organization, and extracellular matrix (ECM) interactions, while mouse models, though valuable, often fall short in representing the unique clinical and biological features of human tumors^18,19^. By contrast, 3D NB models, including multicellular spheroids, bioengineered scaffolds, and microfluidic tumor-on-chip systems, recapitulate the structural and functional complexity of pediatric cancers. These platforms naturally generate gradients of oxygen, nutrients, and drugs, as well as hypoxic, matrix-rich cores that critically shape nanoparticle penetration, retention, and thermal distribution during hyperthermia^18,19^.

Altogether, integrating biomimetic 3D NB models with hyperthermia-based nanotherapies offers a powerful strategy to bridge the translational gap between conventional 2D in vitro assays and in vivo preclinical models. Such platforms not only enable evaluation of MH and PTT under physiologically relevant conditions, but can also provide mechanistic insights into therapy–microenvironment interactions. Ultimately, they can guide the rational optimization of nanoparticle design and treatment parameters, while reducing the number of research animals in preclinical trials tailored for pediatric oncology.

Here, we report a direct comparison of MH and PTT in 3D tissue-engineered NB models based on collagen I/hyaluronic acid scaffolds^20,21^. Using validated iron oxide-based nanoparticles^21^, we assessed heating performance, cell viability, histological markers, and functional outcomes. Our results show that while PTT produced no measurable effects beyond baseline heating, MH triggered a transient antiproliferative response, marked by reduced Ki67 expression and DNA content at 48 h post-treatment. Yet this effect was not sustained: by day 5, proliferation had returned to baseline, and caspase-3/7 analysis revealed no activation of apoptotic signaling. Together, these findings indicate that MH can provide short-term suppression of tumor growth in biomimetic 3D NB models but does not generate lasting therapeutic effects. This highlights the importance of developing rational combination strategies to unlock the full potential of hyperthermia in pediatric oncology, while promoting advanced 3D models as alternatives to minimize animal use in preclinical testing.

## Materials and Methods

### Nanoparticle synthesis and functionalization

Polyacrylic acid–coated magnetite nanoparticles (Fe₃O₄@PAA NPs) were synthesized in aqueous medium using a modified hydrothermal protocol. Briefly, FeCl₂·4H₂O (1.59 g, 8 mmol; Sigma–Aldrich) and FeCl₃·6H₂O (3.78 g, 14 mmol; Sigma–Aldrich) were dissolved in 10 mL of Milli-Q water. Separately, poly(acrylic acid sodium salt) (PAANa; 2.00 g, 0.4 mmol; Sigma–Aldrich) was dissolved in 5 mL of Milli-Q water and combined with the iron solution. Ammonium hydroxide (28–30%; 15 mL; Sigma–Aldrich) was added, producing a black suspension that was transferred to a PTFE vessel and heated to 150 °C for 48 h in a stainless-steel autoclave. After cooling, the product was washed by repeated precipitation in acetone and centrifugation (8,000 rpm, 5 min), resuspended in Milli-Q water, and cleared of large aggregates by low-speed centrifugation (3,000 rpm, 5 min). The resulting stable colloid was stored at 4 °C until use.

For fluorescent labeling, 1 mL of nanoparticle suspension was diluted in 4 mL Milli-Q water and reacted overnight at room temperature with sulforhodamine 101 cadaverine (1 mg in 200 μL DMF; Quimigen) in the presence of excess EDC (Sigma–Aldrich). The product was purified by repeated acetone precipitation and resuspension in Milli-Q water. To further functionalize, glucosamine hydrochloride (50 mg; TCI) was coupled using excess EDC under the same conditions. Final nanoparticles were purified as above, pooled, and stored in Milli-Q water at 4 °C in the dark.

### Scaffold preparation

Porous collagen I/hyaluronic acid (ColI/HA) scaffolds were fabricated by freeze-drying, following previously described protocols^19,20,21^. Briefly, a 1% (w/v) hyaluronic acid (HA) solution was prepared by dissolving 20 mg of sodium hyaluronate (8–15 kDa, Contipro) in 2 mL of distilled water on ice. Collagen I (Corning, #354249) was then mixed with the 1% HA solution at a 4:1 ratio. Aliquots of 50 µL of the ColI/HA suspension were dispensed into 0.5-mL Eppendorf caps used as molds, frozen at –40 °C for 4 h, and lyophilized overnight (–50 °C, 0.12 mbar; Alpha 1-4 LD, Christ). Dry scaffolds were crosslinked in 95% ethanol containing 33 mM EDC and 6 mM NHS (Sigma–Aldrich) for 4 h at 25 °C, washed thoroughly in distilled water (5 × 5 min), re-frozen for 4 h, and lyophilized a second time.

### Cell culture and seeding of scaffolds

Neuroblastoma cell lines SK-N-BE(2), SK-N-LP, SK-N-AS, LA155n and LAN1 were maintained in RPMI-1640 medium (Sigma, R8758) supplemented with 10% heat-inactivated fetal bovine serum (FBS; Gibco) and 1% penicillin/streptomycin at 37 °C in a humidified atmosphere with 5% CO₂.

For scaffold seeding, ColI/HA scaffolds were sterilized in 70% ethanol for 1 h, rinsed, and pre-incubated in RPMI medium for 1 h, as previously described^21,22^. A suspension of 1 × 10⁶ neuroblastoma cells was then loaded onto each scaffold and incubated on a rotary platform at 37 °C for 4 h. Subsequently, scaffolds were transferred to non-treated 24-well plates (Nunc) and cultured in 2 mL of medium to generate tissue-engineered neuroblastoma (TE-NB) models for further studies.

### Nanoparticle treatment of TE-NB models

Tissue-engineered NB (TE-NB) scaffolds were seeded with neuroblastoma cells and maintained in culture for 7 days prior to magnetite nanoparticles (MNPs) treatment. At day 7, constructs were incubated overnight with culture medium containing MNPs (50 µg Fe mL⁻¹) (**Figure 1**). Following incubation, residual nanoparticles were removed by washing, and constructs were placed in fresh medium without MNPs. Samples were subsequently allocated for histology, hyperthermia treatment, or functional assays at defined time points after nanoparticle loading.

**Figure 1.**
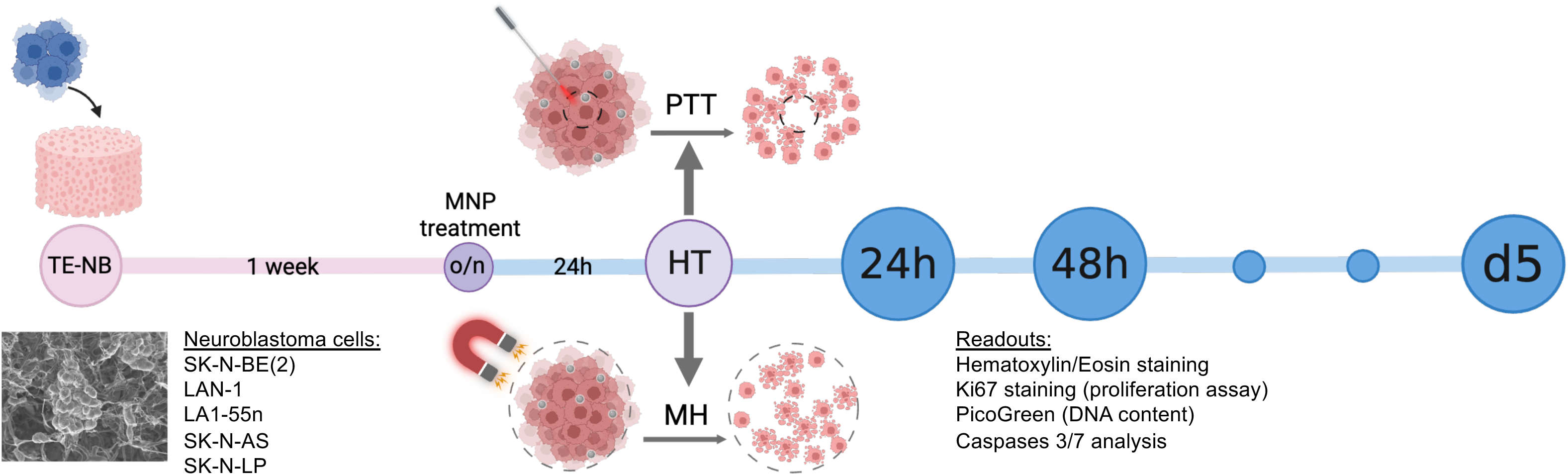
Experimental workflow and timeline of hyperthermia studies in TE-NB models. Schematic overview of the methodology. Tissue-engineered neuroblastoma (TE-NB) scaffolds were cultured for 7 days prior to magnetic nanoparticle (MNP) treatment. Constructs were then incubated overnight with medium containing MNPs (50 µg Fe mL⁻¹), washed to remove residual particles, and maintained in fresh medium without MNPs. Hyperthermia treatments (HT) were applied as either photothermal therapy (PTT; 808-nm NIR laser, 1 W, 10 min) or magnetic hyperthermia (MH; alternating magnetic field, 285 kHz, 20 mT, 60 min). Constructs were harvested at defined time points (24 h, 48 h, and 5 d) post-treatment for assessment of DNA content, cell viability, proliferative activity (Ki67 immunohistochemistry), and apoptotic responses (caspase-3/7 staining).

### Magnetic Resonance Imaging (MRI)

MRI was used to confirm MNP uptake and retention within 3D neuroblastoma scaffolds. Constructs incubated with MNPs (50 µg Fe mL⁻¹) were washed in PBS and fixed in 4% paraformaldehyde. Imaging was performed on a clinical field (3.0 T) horizontal bore MR Solutions Benchtop MRI system, MRS 3000, equipped with 48 G/cm actively shielded gradients. To image the samples, a 56-mm diameter quadrature birdcage coil was used in transmit/receive mode. All MR images were acquired with an image matrix 256 × 252, FOV 60 × 60 mm, 6 slices each with a slice thickness of 1 mm and 0 slice gap. For *T*_2_-weighted imaging, fast spin echo (FSE) sequences with the following parameters were used: T_E_=68 ms, T_R_=4800 ms, N_A_=10, A_T_=24 m 53 s.

### Histology and Prussian blue staining

Scaffolds were washed in PBS and fixed overnight in 10% neutral buffered formalin (Sigma–Aldrich) at 4 °C. After dehydration through ethanol gradients, samples were paraffin-embedded, sectioned (5 μm), and stained with hematoxylin/eosin (H&E).

For iron detection, serial sections were processed with the Hematognost Fe kit (Sigma–Aldrich) according to the manufacturer’s protocol. Iron deposits appeared blue, and nuclei were counterstained with Nuclear Fast Red. Sections were mounted in VectaMount (Vector Labs).

### Hyperthermia treatments

The experimental timeline and analytical pipeline are summarized in **Figure 1**. Briefly, scaffolds were prepared, seeded, and incubated with magnetic nanoparticles (MNPs) for 24 h prior to treatment. After washing to remove unbound particles, constructs were exposed to either photothermal therapy (PTT) or magnetic hyperthermia (MH). PTT was applied using an 808-nm near-infrared laser (Ostech dst11-dilas 50w-808 nm-0.22NA-t19286-v1-503, 1 W, 10 min), while MH was delivered under an alternating magnetic field (NanoTherm, NanoTherics, 285 kHz, 20 mT, 60 min). Control groups included untreated and MNP-only scaffolds. Constructs were collected at defined time points post-treatment (24 h, 48 h, and 120 h) for downstream analyses, including DNA quantification, cell viability, Ki67 immunohistochemistry, and caspase-3/7 activation.

Temperature was monitored using fiber optic probes and recorded simultaneously in the sample of interest and control samples.

### Evaluation of hyperthermia effects

DNA quantification: Cell number in scaffolds was assessed using the Quant-iT PicoGreen dsDNA kit (Life Technologies) following the manufacturer’s instructions.

<u>Viability assay:</u> TE-NB constructs were incubated with 2 μM Calcein AM and counterstained with Hoechst 33342 (16.2 mM; Life Technologies). Images were acquired using an Olympus IX81 fluorescence microscope.

#### Immunohistochemistry for Ki67

Formalin-fixed, paraffin-embedded (FFPE) sections were deparaffinized at 65 °C for 30 min, rehydrated through graded ethanol, and subjected to antigen retrieval in citrate buffer (pH 6.0, microwave, 15 min). Endogenous peroxidase activity was blocked for 10–20 min, and sections were incubated for 1 h at room temperature (RT) with normal horse serum (VECTASTAIN® Elite ABC Kit, Vector Laboratories) to block nonspecific binding. Slides were then incubated overnight at 4 °C in a humidified chamber with rabbit anti-Ki67 antibody (1:1000, Agilent) diluted in PBS.

On the following day, sections were washed with PBS and sequentially incubated with a biotinylated secondary antibody (VECTASTAIN® Elite ABC reagent), following the manufacturer’s instructions. Staining was developed with peroxidase substrate until optimal signal was reached. Slides were counterstained with Hematoxylin QS, dehydrated, and mounted.

For quantification, images were acquired at 20× magnification from both peripheral (edges) and central (core) regions of each scaffold. Ki67⁺ nuclei were counted relative to the total number of hematoxylin-stained nuclei, normalized to untreated controls (set as 1.0), and expressed as relative % of Ki67⁺ cells.

#### Caspase-3/7 Histological Staining and Quantification

FFPE sections of 3D NB scaffolds were deparaffinized at 65 °C for 30 min and rehydrated through graded solvents (xylene, 2 × 5 min; 100%, 90%, 70%, 50%, and 30% ethanol, 2–5 min each). Slides were rinsed in deionized water and maintained moist throughout the procedure.

Sections were incubated with CellEvent™ Caspase-3/7 Green ReadyProbes™ reagent (R37111, Thermo Fisher Scientific), prepared as 1 drop per 0.5 mL PBS. A total of 40–50 µL were applied per section and incubated at 37 °C for 1 h. After rinsing, nuclei were counterstained with Hoechst 33342 (1:5000 in PBS, 10 min, 37 °C), washed to remove excess dye, and mounted in PBS for imaging.

Fluorescence imaging was performed on a Leica AF7000 microscope (20× objective). For each scaffold, four independent regions (S1–S4) were scanned, each comprising 42 contiguous fields (6 × 7). Images were acquired in 8-bit grayscale (0–255).

Quantification was performed in ImageJ (NIH) using a pixel-based approach, as caspase activation produced heterogeneous intracellular intensities precluding cell segmentation. Nuclei were defined by automatic thresholding of the Hoechst (blue) channel to estimate the total number of cells (blue pixels). Caspase activity was quantified in the green channel by applying intensity thresholds to classify signal into three levels: low, medium, and high. The pixel area in each range was measured, and caspase activity was expressed as the ratio of green to blue pixels per image.

### Statistical Analysis

All quantitative results are presented as mean ± SEM, unless otherwise indicated. Comparisons between two groups were performed using Welch’s t-test, while multiple comparisons were analyzed by one-way ANOVA followed by Tukey’s post-hoc test. Differences were considered statistically significant at p < 0.05 (*p < 0.05, **p < 0.01, ***p < 0.001). Graphs and statistical analyses were generated in GraphPad Prism 9 (GraphPad Software, USA)

## Results

To benchmark nanoparticle-mediated heating, first, cuvette assays (3 mL volume) under continuous 808-nm laser irradiation were performed. PBS alone exhibited minimal temperature elevation, whereas PBS supplemented with magnetite nanoparticles (MNPs, 50 µg Fe mL⁻¹) showed a monotonic rise, reaching ΔT ∼3–4 °C within 5 min (**Figure 2A–D**). Equivalent experiments in culture medium (10% FBS) yielded higher baseline heating due to light absorption by proteins, with or without MNPs, though the presence of MNPs slightly increased the net ΔT over 10 min (**Figure 2E–H**).

**Figure 2.**
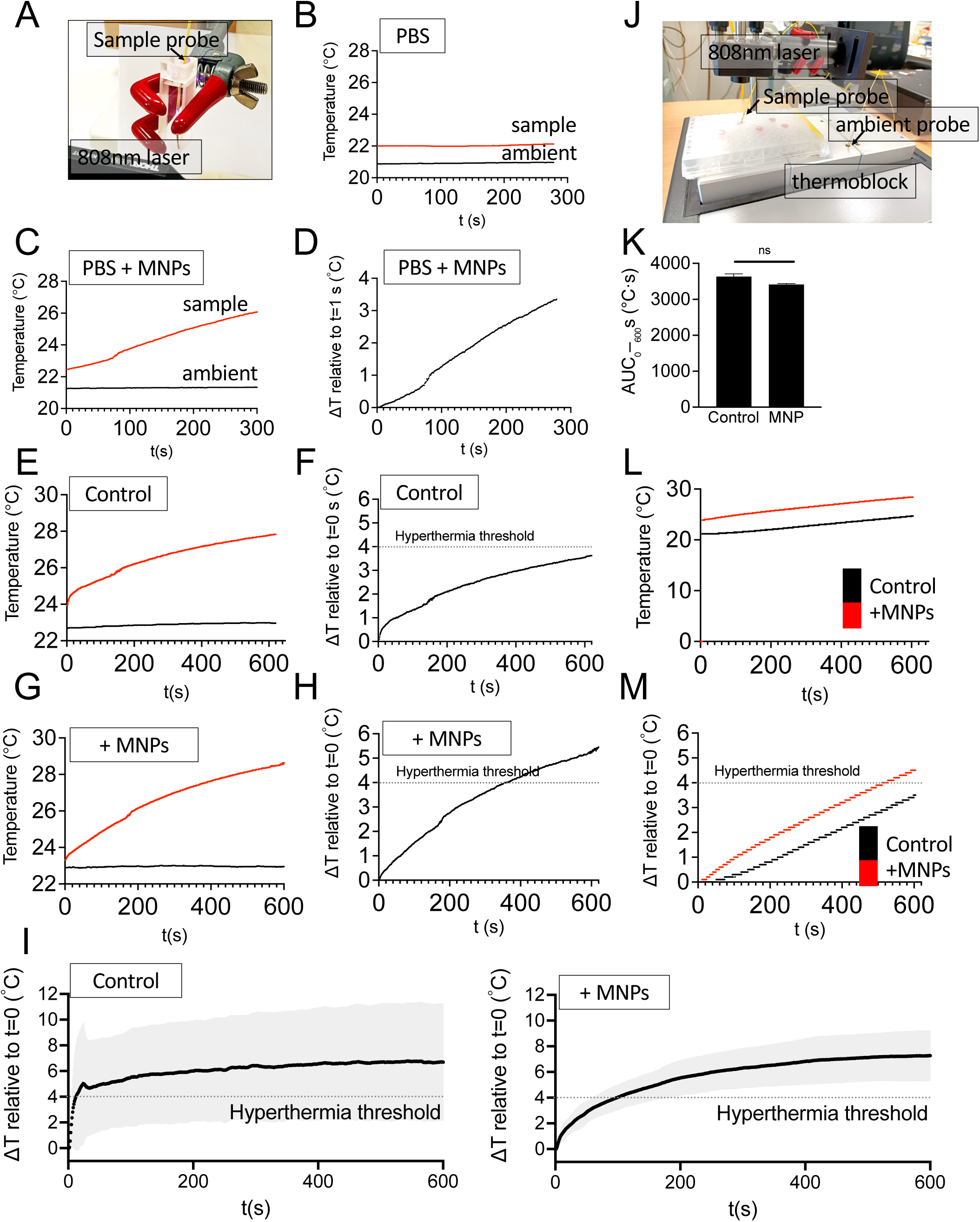
Photothermal and magnetic hyperthermia heating in PBS and culture medium. **(A)** Experimental setup for cuvette assays (3 mL volume; 808-nm diode laser, 1 W; in-sample fiber-optic probe). **(B–C)** Raw temperature traces recorded simultaneously in the irradiated cuvette (“sample”) and the ambient probe for PBS alone (B) and PBS supplemented with magnetite nanoparticles (MNPs, 50 µg Fe mL⁻¹) (C). **(D)** Corrected temperature rise, defined as ΔT = (Tsample − Tambient) relative to the first recorded second, for PBS + MNPs. **(E–H)** Equivalent assays performed in 3-mL cuvettes containing culture medium with 10% FBS with or without MNPs show monotonic heating in both cases, with MNPs reaching slightly higher ΔT over 10 min. (F-H) The dashed line indicates ΔT = 4 °C, corresponding to the hyperthermia threshold (∼41 °C if starting at 37 °C). **(I)** Corrected ΔT versus time in the 96-well plate geometry (200 µL/well culture medium with 10% FBS, mean ± SD, n = 5) with the plate maintained on a 37 °C thermoblock to approximate physiological baseline. The dashed line indicates ΔT = 4 °C, corresponding to the hyperthermia threshold (∼41 °C if starting at 37 °C). Both conditions approached plateaus of ∼7–8 °C above baseline, demonstrating that bulk heating of the medium/plate can reach the therapeutic window even without MNPs. **(J)** Experimental setup for the 96-well plate configuration using a thermoblock. **(K)** Integrated thermal dose (AUC₀–₆₀₀s, °C·s) calculated as the area under the ΔT(t) curve from 0 to 600 s. Bars represent mean ± SEM values for medium with or without MNPs; equivalent values in °C·min are provided in the Results. The integrated thermal load was modestly higher with MNPs but not statistically significant (Welch’s t-test, p > 0.05). **(L)** Raw temperature traces during magnetic hyperthermia in culture medium with or without MNPs under an alternating magnetic field (AMF; 285 kHz, 20 mT). **(M)** Corrected ΔT traces under AMF exposure show that samples containing MNPs exhibited a clear monotonic increase in temperature, reaching ΔT ∼4 °C within 10 min, whereas medium alone showed negligible heating under identical conditions.

Next, heating in a 96-well plate format (200 µL/well, **Figure 2J**) maintained at 37 °C to approximate physiological baseline was evaluated. Both medium alone and medium + MNPs approached plateaus of ∼7–8 °C above baseline, surpassing the hyperthermia threshold (∼42 °C). However, integrated thermal dose analysis (AUC₀–₆₀₀s) revealed no significant difference between conditions (Welch’s t-test, p > 0.05), indicating that bulk medium absorption dominates thermal load in this geometry (**Figure 2I–K**).

Finally, under alternating magnetic fields (AMF; 285 kHz, 20 mT), only wells containing MNPs displayed measurable heating, reaching ΔT ∼4 °C within 10 min, whereas medium alone showed negligible response (**Figure 2L–M**). These data confirm that, while photothermal heating in vitro is largely governed by nonspecific medium absorption, AMF exposure produces MNP-dependent heating with minimal background, validating the system for subsequent 3D experiments.

To establish the experimental workflow, tissue-engineered NB (TE-NB) scaffolds were generated first by culturing cells for 7 days prior to MNP treatment (**Figure 1**). At day 7, constructs were incubated overnight with medium containing MNPs (50 µg Fe mL⁻¹), washed to remove residual particles, and maintained in fresh medium without MNPs, as previously described^21^.

To evaluate short-term cytocompatibility, Calcein AM/Hoechst staining of SK-N-BE(2) scaffolds at 24 h post–MNP treatment showed uniformly high viability in both control and MNP-loaded conditions, with no evidence of acute cytotoxicity (**Figure 3A**). Consistently, DNA quantification by PicoGreen revealed no significant difference in cell number between groups at 24 h (**Figure 3B**).

**Figure 3.**
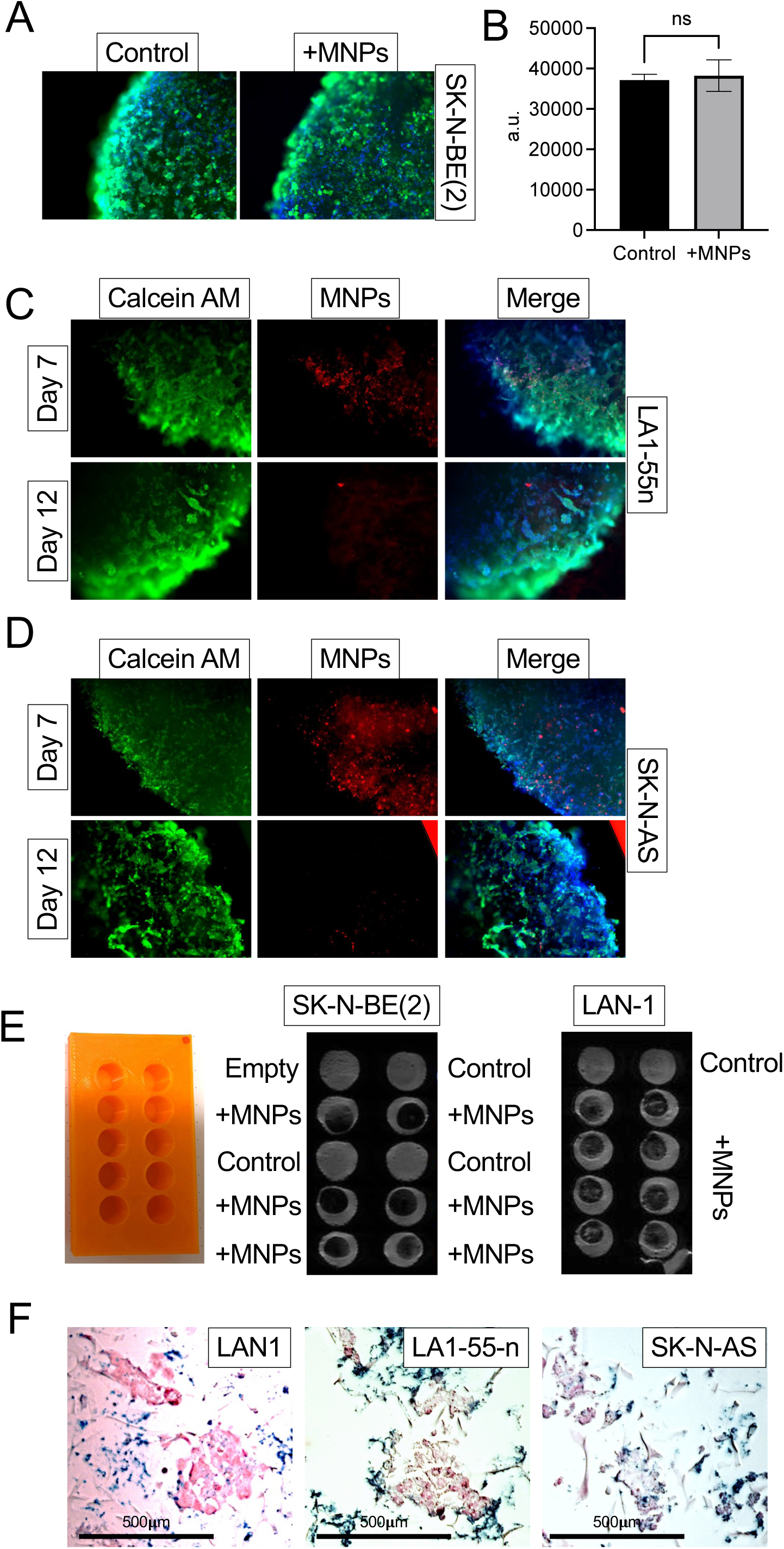
Uptake and biocompatibility of magnetic nanoparticles (MNPs) in 3D neuroblastoma scaffolds. **(A)** Calcein AM staining of SK-N-BE(2) scaffolds at 24 h post-treatment in the absence (control) or presence of MNPs. Green = calcein-positive viable cells; blue = Hoechst 33342, nuclei. **(B)** DNA quantification by PicoGreen at 24 h in SK-N-BE(2) scaffolds with or without MNPs. Bars represent mean ± SEM values (n = 4); ns = not significant (Welch’s t-test). **(C–D)** Fluorescence microscopy of LA1-55n (D) and SK-N-AS (E) scaffolds at 7 and 12 days after MNP treatment. Images show viable cells (green, calcein), nuclei (blue, Hoechst 33342), and intracellular MNPs (red puncta). Scale bars, 100 µm. **(E)** Magnetic resonance imaging (MRI) at 48 h of collagen I–hyaluronic acid scaffolds seeded with SK-N-BE(2) and LAN-1 cells after MNP treatment. Left: Custom 3D-printed PLA holder used for positioning the samples. Right: representative *T_2_*-weighted images. **(F)** Prussian blue staining of SK-N-BE(2), LA1-55n, and SK-N-AS scaffolds at 48 hours after MNP treatment. Blue deposits indicate intracellular iron; nuclei are counterstained with nuclear fast red (pink/red). Scale bars, 500 µm. Representative images are shown (n = 4).

Next, nanoparticle persistence and distribution across multiple NB lines was investigated. Fluorescence imaging confirmed intracellular MNPs (red puncta) co-localizing with viable cells (green, Calcein AM) and nuclei (blue, Hoechst 33342) in LA1-55n and SK-N-AS scaffolds at day 7 (**Figure 3C–D**). While MNPs remained readily detectable at this time point, their signal was markedly reduced by day 12 post–MNP treatment, suggesting progressive redistribution, dilution during cell proliferation, or clearance processes, consistent with prior reports in TE-NB using SK-N-BE(2) cells^21^.

Complementary non-invasive medical imaging approaches further validated nanoparticle uptake and retention. At a scaffold level, MRI scans performed at 48 h confirmed detectable iron-associated signal destruction in *T_2_*-weighted images within collagen I–hyaluronic acid scaffolds seeded with SK-N-BE(2) and LAN-1 cells (**Figure 3E**). In parallel, at a cellular/sub-cellular level, histological Prussian blue staining at 48 h post–MNP treatment verified intracellular iron deposition across three NB cell lines (SK-N-BE(2), LA1-55n, SK-N-AS), with blue iron deposits observed in the cytoplasm and nuclei counterstained in red (**Figure 3F**). Together, these results demonstrate that MNPs can be efficiently internalized and retained by multiple NB cell lines in 3D scaffolds without acute cytotoxic effects, confirming their suitability for subsequent hyperthermia experiments.

To evaluate the short-term impact of hyperthermia on proliferation, TE-NB constructs from five NB cell lines were treated either with PTT, where laser energy was directed to the scaffold core, or with MH, where the alternating magnetic field uniformly encompassed the entire construct. Ki67 immunohistochemistry was performed at 24 h post-treatment to assess proliferative responses across edge and inner regions. Both modalities produced heterogeneous outcomes depending on the cell line and scaffold region, with changes manifesting as either increases or decreases in Ki67 labeling relative to untreated controls (**Figure 4**).

**Figure 4.**
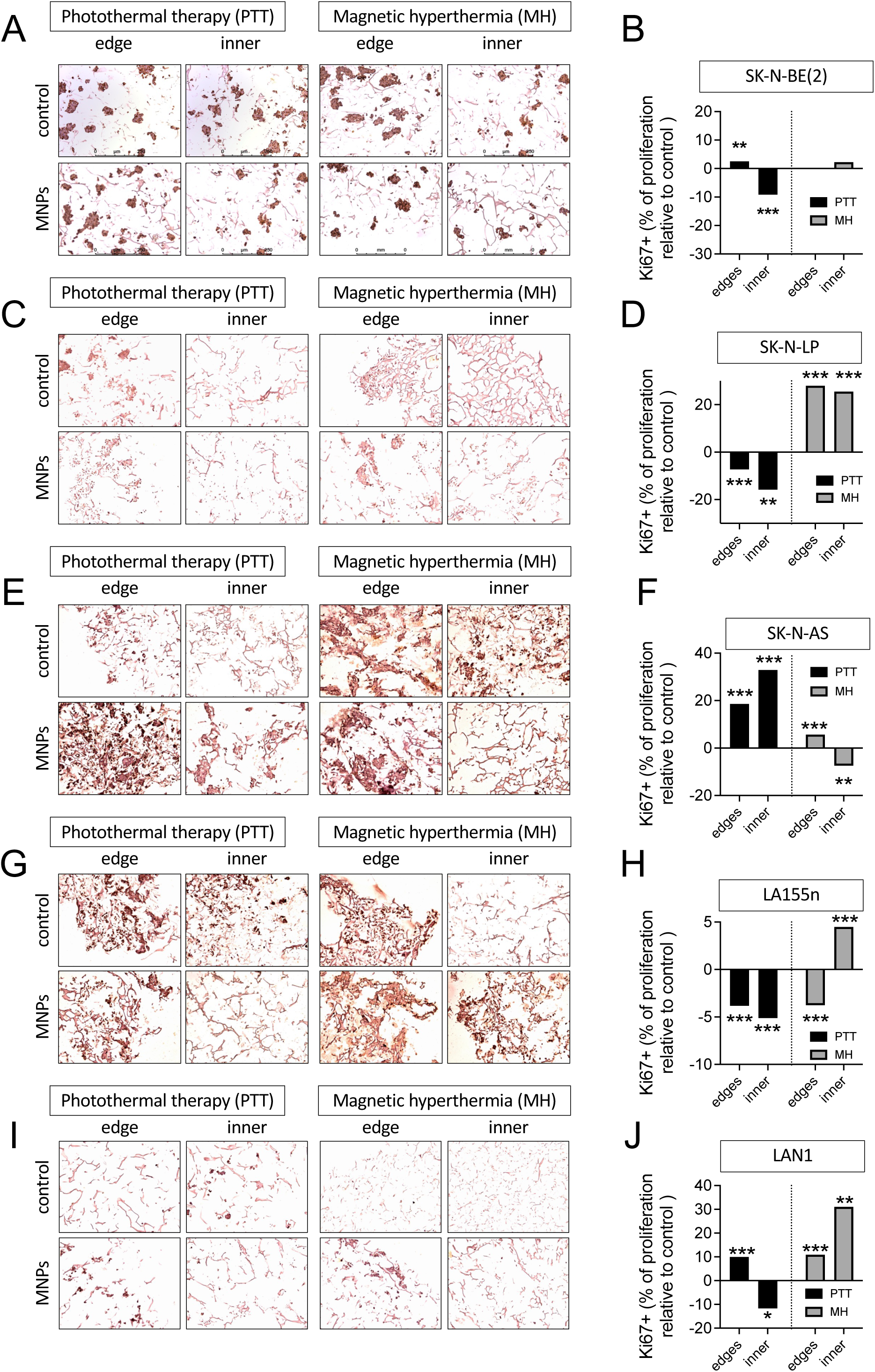
Effect of photothermal therapy (PTT) and magnetic hyperthermia (MH) on tissue-engineered neuroblastoma (TE-NB) models at 24 h post-treatment. (A, C, E, G, **I)** Representative Ki67 immunohistochemistry in TE-NB constructs derived from five neuroblastoma cell lines: SK-N-BE(2), SK-N-LP, SK-N-AS, LA1-55n, and LAN-1, respectively. Columns depict edge and inner scaffold regions, with untreated controls and MNP-treated samples shown as indicated. (**B, D, F, H, J**) Quantification of Ki67-positive nuclei (% relative to untreated controls) for each corresponding model, separated by edge and inner compartments. Bars represent mean ± SEM; individual data points are shown where available. Statistical significance versus untreated controls: *p < 0.05, **p < 0.01, ***p < 0.001. Scale bars as indicated.

In SK-N-BE(2) constructs, PTT significantly increased Ki67 labeling at the edges but reduced it in the inner regions compared to controls, whereas MH did not induce significant differences (**Figure 4A–B**). In SK-N-LP scaffolds, PTT led to a significant reduction of Ki67 in both edges and inner cores, while MH produced a significant increase in both regions (**Figure 4C–D**). SK-N-AS constructs showed a significant increase in Ki67 labeling with PTT across both regions; under MH, proliferation increased at the edges but decreased in the inner compartment (**Figure 4E–F**). In LA1-55n models, PTT reduced Ki67 labeling in both edges and inner regions, whereas MH decreased proliferation at the edges but enhanced it in the inner regions (**Figure 4G–H**). Finally, in LAN-1 scaffolds, PTT increased Ki67 staining at the edges but reduced it in the inner regions; conversely, MH elevated proliferation at both edges and inner cores compared with controls (**Figure 4I–J**).

Together, these findings demonstrate that both PTT and MH can modulate proliferation in 3D NB models within 24 h, but the direction and magnitude of the effect vary markedly by cell line and scaffold region. This heterogeneity highlights the importance of evaluating multiple NB subtypes and spatial compartments when assessing hyperthermia responses in biomimetic models.

To extend the analysis to later time points, TE-NB constructs based on SK-N-BE(2) cells were evaluated at 48 h after hyperthermia. H&E staining showed that overall scaffold architecture and cell density were preserved in control and PTT-treated samples, whereas MH-treated constructs displayed clear signs of structural disruption and cell loss (**Figure 5A**). In line with these observations, Ki67 immunohistochemistry revealed divergent effects of the two modalities: PTT induced a significant increase in proliferative labeling in both edge and inner regions, while MH caused a marked reduction across the same compartments (**Figure 5B–C**). Quantitative DNA analysis by PicoGreen further supported these findings, with a significant decrease in DNA content detected only in MH-treated constructs, indicating treatment-induced cell loss, whereas PTT had no measurable impact compared with controls (**Figure 5D**). Together, these results demonstrate that MH exerts a more pronounced cytotoxic effect on SK-N-BE(2) TE-NB models than PTT under the applied parameters.

**Figure 5.**
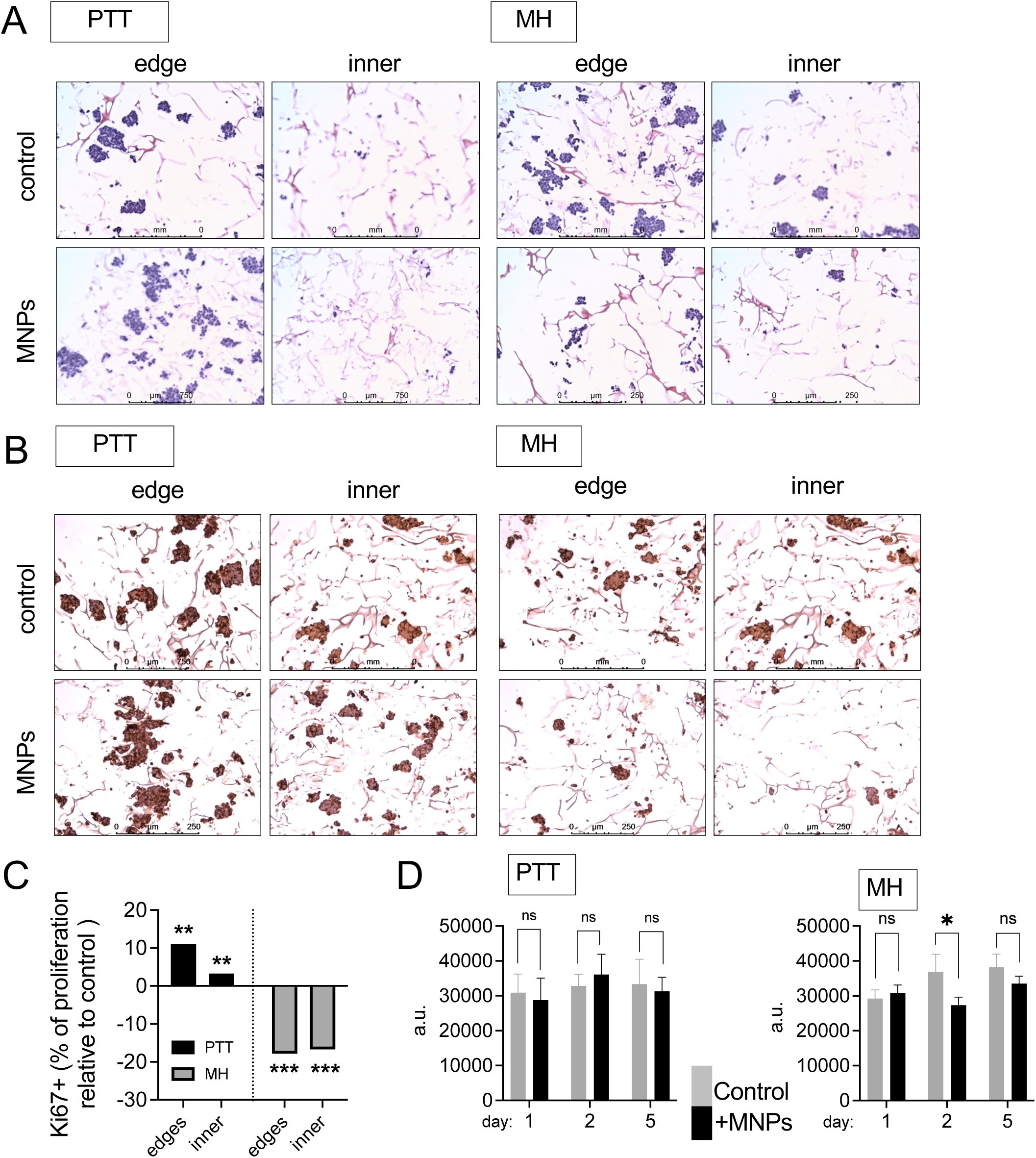
Effect of photothermal therapy (PTT) and magnetic hyperthermia (MH) on tissue-engineered neuroblastoma (TE-NB) models at 48 h post-treatment. **(A)** Representative H&E staining of TE-NB constructs. Control and PTT-treated samples preserved their overall architecture and cell density, whereas MH-treated constructs exhibited marked tissue disruption and cell loss by 48 h. **(B)** Ki67 immunohistochemistry in SK-N-BE(2) constructs at 48 h post-treatment, showing edge and inner regions in control, PTT-, and MH-treated samples. **(C)** Quantification of Ki67-positive cells in SK-N-BE(2) constructs. PTT induced a significant increase in proliferative labeling in both edge and inner regions compared to controls, whereas MH caused a pronounced reduction in Ki67 expression in both regions. **(D)** DNA quantification by PicoGreen assay in SK-N-BE(2) constructs at 48 h. MH treatment significantly reduced DNA content (p < 0.05 vs. control), consistent with cell loss, while PTT-treated constructs showed no significant changes. Data represent mean ± SD.

To assess apoptosis after photothermal therapy, caspase-3/7 activation was evaluated in TE-NB constructs at 24 h, 48 h, and 120 h (5 days) post-PTT using a pixel-based quantification of CellEvent™ staining on FFPE sections. Representative confocal images did not reveal qualitative increases in green caspase signal in PTT-treated samples compared with untreated controls at any time point (**Figure 6A**). Quantitatively, the green-to-blue pixel ratio remained comparable between groups at 24 h, 48 h, and 120 h, with no statistically significant differences detected **(Figure 6B**). Stratification of the green channel into low, medium, and high intensity bands yielded similar results, with no time-or dose-dependent increases in caspase-3/7 activity in PTT-treated constructs relative to controls. These data indicate that, under the applied PTT parameters, apoptosis was not detectable in TE-NB scaffolds up to 5 days post-treatment.

**Figure 6.**
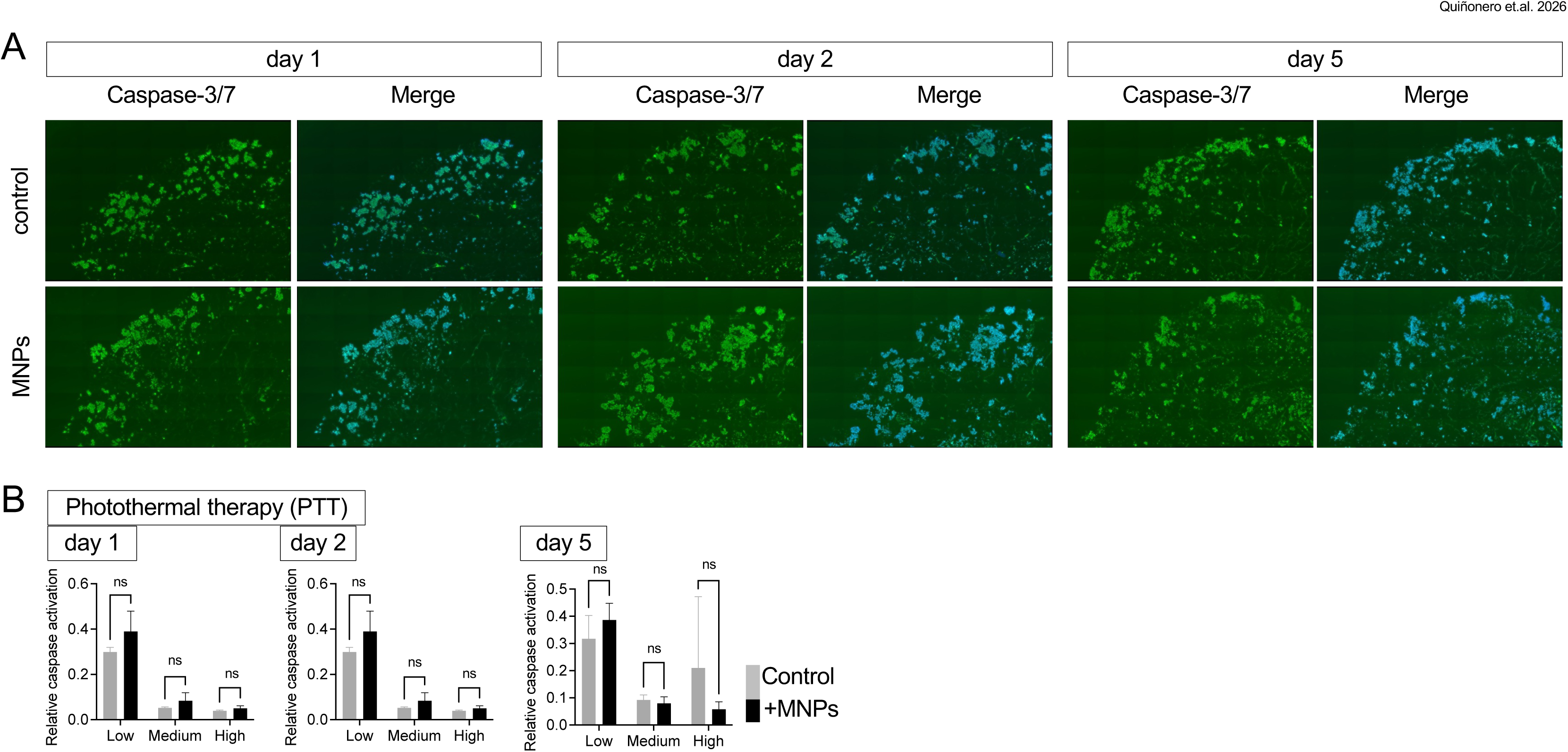
Caspase-3/7 activation following photothermal therapy (PTT) in tissue-engineered neuroblastoma (TE-NB) models. **(A)** Representative confocal images of CellEvent™ Caspase-3/7 (green) with nuclear counterstain (blue, Hoechst 33342) acquired at 24 h (Day 1), 48 h (Day 2), and 120 h (Day 5) after PTT. Untreated controls and PTT-treated constructs (MNP-loaded) are shown for each time point. **(B)** Quantification of caspase-3/7 activity using a pixel-based approach (green/blue pixel ratio per image) across 24 h, 48 h, and 120 h post-PTT. Distributions are also reported by intensity bands (low/medium/high). Bars represent mean ± SD; no significant differences (ns) were detected between PTT-treated and control groups at any time point or intensity band (n as indicated; statistical tests in Methods)

Likewise, to investigate whether magnetic hyperthermia induces apoptotic signaling, caspase-3/7 activation was analyzed in TE-NB constructs up to 5 days post-treatment. Confocal imaging of CellEvent™ staining revealed no obvious qualitative increase in caspase signal in MH-treated constructs compared with controls at 24 h, 48 h, or 120 h (5 days) **(Figure 7A)**. Quantitative pixel-based analysis confirmed the absence of significant treatment-related differences at any time point, with the exception of a minor shift detected at 24 hours in the high-intensity band, where control scaffolds displayed a greater caspase signal than MH-treated constructs **(Figure 7B).** These findings indicate that MH, despite its pronounced effects on proliferation and DNA content (Figure 4), does not elicit detectable caspase-3/7 activation in TE-NB scaffolds within the first 5 days, consistent with a non-apoptotic mechanism of cytotoxicity.

**Figure 7.**
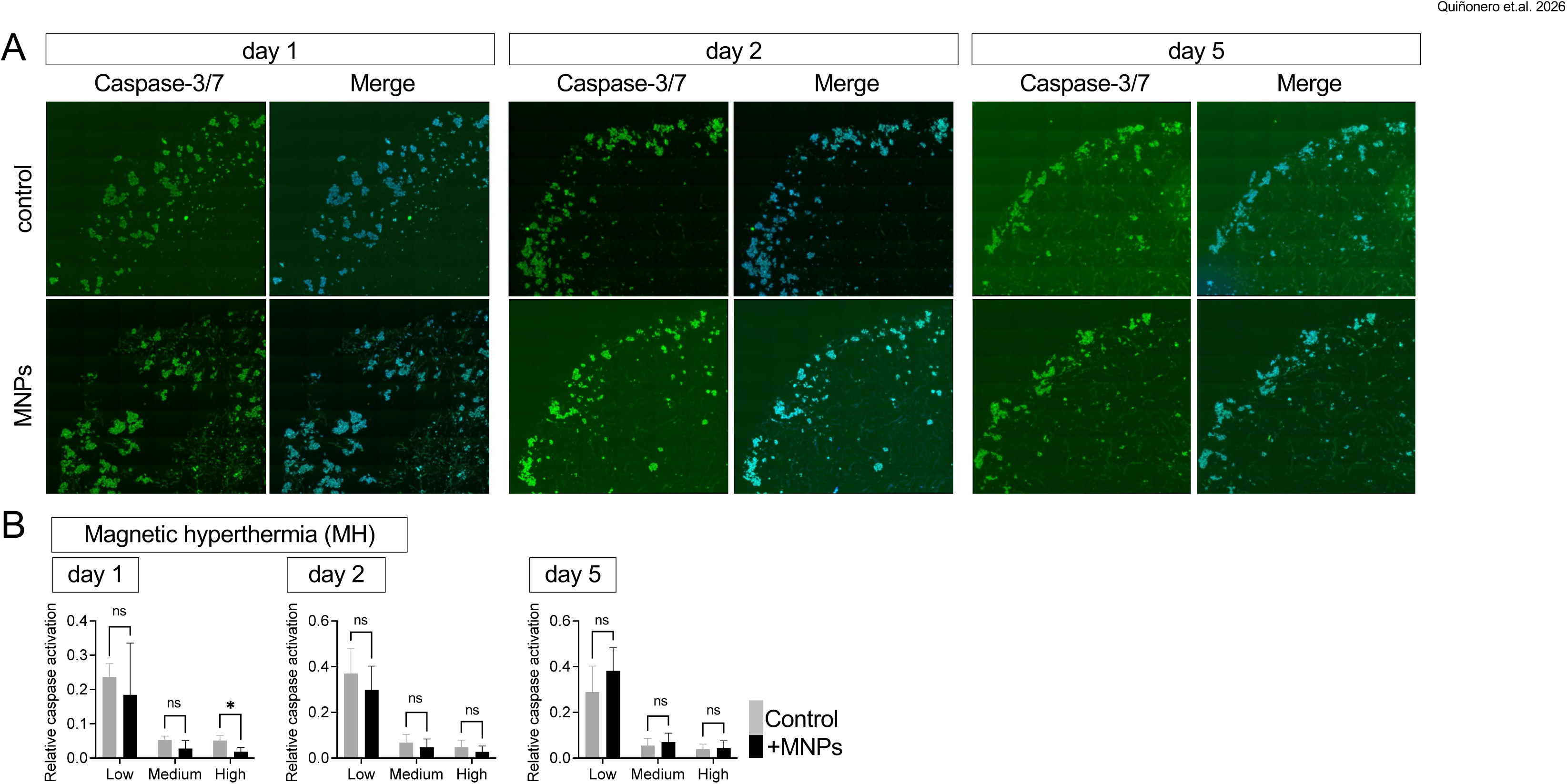
Caspase-3/7 activation following magnetic hyperthermia (MH) in tissue-engineered neuroblastoma (TE-NB) models. **(A)** Representative confocal images of CellEvent™ Caspase-3/7 (green) with nuclear counterstain (blue, Hoechst 33342) acquired at 24 h (Day 1), 48 h (Day 2), and 120 h (Day 5) after MH. Untreated controls and MH-treated constructs (MNP-loaded) are shown for each time point. **(B)** Quantification of caspase-3/7 activity (green/blue pixel ratio per image) across 24 h, 48 h, and 120 h post-MH. Distributions are also reported by intensity bands (low/medium/high). Bars represent mean ± SD. no significant differences (ns) were detected between groups at any time point, except for a modest increase in the “high” intensity fraction at Day 1 in control constructs relative to MH-treated scaffolds (*p < 0.05; n as indicated; statistical tests in Methods).

## Discussion

In this study, the potential therapeutic effects of photothermal therapy (PTT) and magnetic hyperthermia (MH) were compared in 3D tissue-engineered neuroblastoma (TE-NB) models, analyzing heating performance, nanoparticle uptake, proliferative dynamics, and apoptotic responses. Several key insights emerge.

Initial thermal characterization showed that in vitro photothermal heating is largely dominated by nonspecific absorption of culture medium, limiting the contribution of nanoparticles. In contrast, MH generated MNP-dependent heating with minimal background, validating its specificity and supporting its use in 3D tumor models. Uptake and persistence studies further confirmed that MNPs are efficiently internalized across multiple NB lines and retained for several days without acute toxicity, establishing their suitability as functional mediators of hyperthermia.

Proliferation assays revealed that both PTT and MH modulate tumor cell growth within 24 h, but the direction and magnitude of effects varied sharply by cell line and by scaffold region (edge vs. core). This heterogeneity underscores two principles: (i) therapy outcomes are strongly shaped by microenvironmental gradients (nutrients, oxygen, nanoparticle distribution, heat dissipation), and (ii) neuroblastoma subtypes display intrinsic variability that dictates response. A single thermal input could trigger either suppression or paradoxical stimulation of proliferation, highlighting the limitations of 2D for predicting efficacy.

At 48 h, SK-N-BE(2) constructs revealed a clear divergence between modalities. PTT not only failed to suppress proliferation but was associated with increased Ki67 labeling, consistent with a stress-induced rebound. MH, by contrast, disrupted scaffold architecture, reduced DNA content, and suppressed proliferation across regions, confirming its stronger cytotoxic effect. These results highlight a key advantage of MH: homogeneous field penetration provides more consistent thermal distribution compared with the spatial variability of laser-based PTT.

Mechanistically, neither PTT nor MH induced detectable caspase-3/7 activation up to 5 days post-treatment. The absence of apoptotic signatures, despite MH-induced suppression of proliferation and DNA content, suggests that cytotoxicity proceeds through non-apoptotic routes such as necrosis, ferroptosis, metabolic collapse, senescence-like arrest, or, less likely, nanoparticle-driven mechanical disruption^23,24^. This distinction is important: failure to engage apoptosis may allow eventual proliferative recovery, while non-apoptotic death pathways could generate inflammatory signals that either promote immune clearance or favor _relapse23,25,26._

These findings must also be understood through the lens of thermal dose. Hyperthermia spans a biological spectrum ranging from mild hyperthermia (40–43 °C), which perturbs protein folding, redox balance, and metabolic homeostasis, to high-temperature thermal ablation (>50 °C), which produces immediate coagulative necrosis^27^. The temperatures achieved in our scaffold-based system fall within the mild hyperthermia range. In this regime, therapeutic outcomes depend not only on peak temperature but also on exposure time, spatial thermal gradients, and the stress-response thresholds inherent to each NB subtype. This framework helps explain why MH produced progressive cytostasis and architectural disruption that became more evident at 48 h, whereas PTT induced only transient, heterogeneous responses dominated by adaptive proliferation. Subthreshold heating can activate prosurvival pathways, while insufficient thermal gradients may prevent effective cytotoxicity. Incorporating these thermobiological principles is therefore essential for rational optimization of hyperthermia protocols in biomimetic tumor models.

Taken together, these results illustrate both the promise and limitations of hyperthermia in pediatric oncology. MH showed superior ability to suppress proliferation and compromise scaffold integrity, but without durable apoptotic engagement its standalone efficacy is limited. This argues for combinatorial approaches pairing MH with DNA-damaging agents, p53 activators, ferroptosis inducers, or immunotherapies to transform transient cytostasis into lasting tumor control^28–31^. Importantly, non-apoptotic stress may also prime tumors for immune recognition by releasing damage-associated molecular patterns (DAMPs), suggesting a role for MH as an immunogenic adjuvant^10,25,26^.

The paradoxical proliferative surges observed with PTT highlight the risks of suboptimal dosimetry. Sublethal laser exposure may activate stress pathways such as HSF1–HSP90 or PI3K–AKT, transiently enhancing survival and growth^32–34^. MH, though more reliable, also produced variable outcomes across NB lines, reinforcing that hyperthermia should not be seen as a universally cytotoxic modality, but as a context-dependent modulator of tumor fate.

Finally, these findings advance the concept of “precision hyperthermia” as a therapeutic strategy for pediatric oncology. Neuroblastoma exhibits pronounced inter- and intra-patient heterogeneity, arising from variations in microenvironmental architecture, cellular states, and stress-response programs. Such factors critically determine heat propagation, nanoparticle dynamics, and downstream biological outcomes. Consequently, hyperthermia should not be regarded as a uniform intervention but rather tailored to the specific microenvironmental context of each tumor. Our TE-NB platform provides a tractable and patient-relevant system to capture this complexity. By reconstructing scaffolds with patient-derived neuroblastoma cells, and ideally incorporating stromal and immune components, ex vivo dose–response assays could be performed to map individualized thermal sensitivities. These experiments would not only determine optimal therapeutic windows for PTT and MH but also identify rational combinations with chemotherapy, targeted drugs, or immunotherapies tailored to each tumor subtype or patient.

Such an approach directly aligns with the principles of precision medicine: moving away from one-size-fits-all interventions and toward personalized treatment strategies informed by predictive human-based models^35,36^. Importantly, it would reduce reliance on animal models, meeting the ethical and regulatory demands for alternatives in pediatric cancer research. By generating actionable patient-specific data, TE-NB scaffolds could serve as decision-support tools, accelerating the translation of hyperthermia-based regimes into the clinic while ensuring their safety, effectiveness, and biological relevance.

## Conclusion

This study provides the first systematic comparison of photothermal therapy (PTT) and magnetic hyperthermia (MH) in tissue-engineered neuroblastoma (TE-NB) models. By integrating 3D scaffolds with multiple NB cell lines and magnetic nanoparticles, we demonstrated that MH achieves more homogeneous heating and greater suppression of proliferation than PTT, although neither modality induced sustained apoptotic responses. These results indicate that hyperthermia acts primarily through non-apoptotic pathways, with outcomes shaped by tumor subtype and microenvironmental gradients. Rather than a binary cytotoxic agent, hyperthermia should be viewed as a precision modulator of tumor fate. TE-NB platforms provide a powerful route to define individualized therapeutic windows and rational combinations, accelerating translation of hyperthermia-based strategies into pediatric oncology.

## Disclosure

The authors report no conflicts of interest in this work

## Author contributions

GQ, APM and LDG performed experiments and contributed to data analysis and interpretation. JG contributed to conceptualization, performed experiments, analyzed and interpreted results, and edited the manuscript. JM and JS reviewed and edited the manuscript. AV conceived the project, performed experiments, analyzed and interpreted results, and wrote the manuscript.

## Availability of data and materials

The datasets during and/or analyzed during the current study are available from the corresponding author upon reasonable request.

## Acknowledgments

The IBEC Group is supported by Department of Research and Universities of the Generalitat de Catalunya (2021 SGR 01545), the CERCA Programme / Generalitat de Catalunya and the Networking Biomedical Research Center (CIBER) of Spain. CIBER is an initiative funded by the VI National R&D&i Plan 2008-2011, Iniciativa Ingenio 2010, Consolider Program, CIBER Actions and the Instituto de Salud Carlos III (RD16/0006/0012), with the support of the European Regional Development Fund (ERDF); GQ is supported by the Neuroblastoma Foundation; AV is supported by the grant (INVES234968VILL) from the Scientific Foundation of the Spanish Association Against Cancer (AECC), Grant PID2023-153218OB-I00 funded by MICIU/AEI/ 10.13039/501100011033 and by “ERDF/EU”, grant PID2020-117977RA-I00 by AEI, and the Association of Families and Friends of Children with Neuroblastoma (NEN); APM acknowledges support from the European Union’s Horizon Europe research and innovation programme under the Marie Skłodowska-Curie Actions Postdoctoral Fellowship grant agreement No 101149210.

